# Bacterial physiological adaptations to contrasting edaphic conditions identified using landscape scale metagenomics

**DOI:** 10.1101/117887

**Authors:** Ashish A. Malik, Bruce C. Thomson, Andrew S. Whiteley, Mark Bailey, Robert I. Griffiths

## Abstract

Environmental factors relating to soil pH are widely known to be important in structuring soil bacterial communities, yet the relationship between taxonomic community composition and functional diversity remains to be determined. Here, we analyze geographically distributed soils spanning a wide pH gradient and assess the functional gene capacity within those communities using whole genome metagenomics. Low pH soils consistently had fewer taxa (lower alpha and gamma diversity), but only marginal reductions in functional alpha diversity and equivalent functional gamma diversity. However, coherent changes in the relative abundances of annotated genes between pH classes were identified; with functional profiles clustering according to pH independent of geography. Differences in gene abundances were found to reflect survival and nutrient acquisition strategies, with organic-rich acidic soils harboring a greater abundance of cation efflux pumps, C and N direct fixation systems and fermentation pathways indicative of anaerobiosis. Conversely, high pH soils possessed more direct transporter-mediated mechanisms for organic C and N substrate acquisition. These findings show that bacterial functional versatility may not be constrained by taxonomy, and we further identify the range of physiological adaptations required to exist in soils of varying nutrient availability and edaphic conditions.

## Introduction

Understanding the key drivers and distributions of microbial biodiversity from both taxonomic and functional perspectives is critical to understand element cycling processes under different land management and geo-climatic scenarios. Distributed soil surveys have shown strong effects of soil properties on the taxonomic biodiversity of bacterial communities (1–5), and to a lesser degree for other soil microbes such as the fungi and protozoa (6, 7). Particularly for bacteria, soil pH often appears as a strong single correlate of biodiversity patterns. This is either due to the direct effects of acidity, or soil pH representing a proxy for a variety of other factors across soil environmental gradients. Acidic soils generally harbor reduced phylogenetic diversity and are usually dominated by acidophilic Acidobacterial lineages. Intermediate pH soils (pH 5-7) are generally composed of larger numbers of taxa, often with a few dominant lineages; whereas at neutral pH (typically intensive agricultural soils) a more even distribution of a multitude of taxa is typically observed (1, 2, 8). A fundamental issue to be resolved is whether these differences in taxonomic diversity reflect changes in functional genetic potential. Many dominating organisms, particularly in the more oligotrophic habitats are difficult to culture, so we know little of the functional characteristics of these organisms at either the phenotypic or genomic level. These knowledge gaps ultimately limit the utility of taxonomic methods (e.g rRNA based) for further understanding ecosystem function.

Whole genome shotgun sequencing has been applied to elucidate the functional diversity of communities from a range of ecosystems (8–12), and its application provides novel insights into the genetic diversity of biochemical processes occurring in soils as well as permitting ecological investigations of microbes within a functional trait based context. The established relationships between edaphic conditions and taxonomic biodiversity can now be complemented with a concurrent understanding of altered environmental microbial physiology and metabolism, which can provide better understanding of the cycling of elements in soils. In the first instance, we need to identify the range of functional genes present in soils to better elucidate the genetic determinants of relevant environmental processes (for example determining the dominant biochemical pathways most likely to be contributing to fluxes). Second, we need to identify the ecological mechanisms determining how genetic pathways are altered according to environmental change. Addressing these knowledge gaps will lead to a better understanding of how will bacterial diversity relates to functional potential, and importantly will elucidate how environmental change can impair or enhance soil functionality.

Here we seek to test whether known differences in taxonomic diversity and composition are reflected in functional gene profiles, by implementing metagenomic sequencing of geographically dispersed soils at opposing ends of a temperate bacterial diversity gradient. We chose to sequence 4 low and 4 high pH soils which had previously been collected as part of a national survey of Britain (figure 1) and were known to comprise low and high taxonomic diversity respectively (7). In addition to assessing richness effects, we seek to explore change in specific functional gene abundances in order to elucidate the physiological constraints acting on different soil systems and identify variance in functional pathways of relevance to soil biogeochemical cycling.

**Figure 1:**
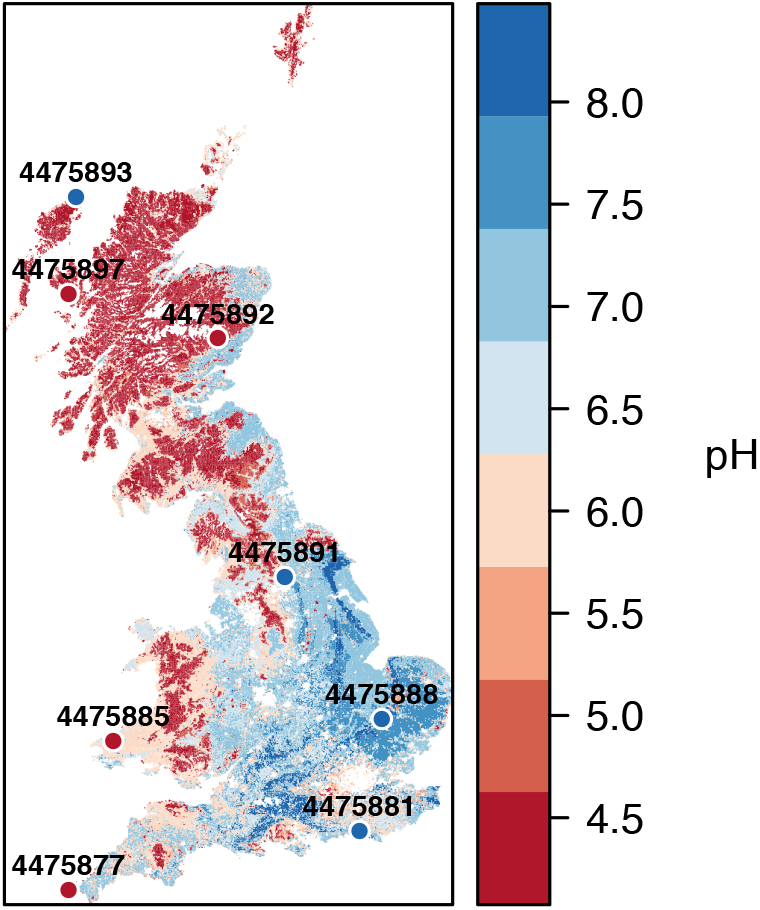
Geographically distributed soils from a range of habitats sampled at opposing ends of a landscape pH gradient. The sampling locations are displayed over a soil pH map of Britain derived from the UK Soils portal (http://ukso.org)

## Results and discussion

### Differences in taxonomic richness are not reflected in functional richness

Four geographically distributed soils exhibiting similar pH were selected to represent each of the low and high pH classes based on our previous work. These encompassed a variety of different habitats (table 1), with the low pH soils typically being found at higher altitude and possessing higher organic matter and moisture content consistent with broader patterns across Britain. Amplicon sequencing confirmed that the microbial taxonomic richness differed between the two soil pH groups, with both alpha and gamma diversity being higher in the high pH soil communities (figure 2a). We then sought to examine whether the richness of annotated genes from metagenomic sequencing was also reduced in low pH soils. Metagenomic sequencing using the Roche 454 platform resulted in a total of 4.9 million quality filtered reads across all samples; with 0.4 to 0.7 million reads per sample (table 1). An average of 8.3±0.4 % of sequences failed to pass the MG-RAST QC pipeline. Following gene annotation using MG-RAST using default stringency criteria, the percentage of reads annotated to predicted proteins ranged from 63.3 to 73.7 % across samples (average=68.9 %). The majority of annotations were assigned to bacterial taxa (94.7±1.6% in low pH soils, 96.6±0.3% in high pH soils); with only a small proportion being eukaryotic (3±1.2% in low pH soils, 1.8±0.1% in high pH soils) and archaeal (2.2±1.4% in low pH soils, 1.4±0.3% in high pH soils). The marked low proportion of fungi contributing to the soil DNA pool was also reflected in the RDP annotation of ribosomal RNA reads (not shown). The metagenomic guanine plus cytosine content (GC%) was lower in acidic anaerobic soils (table 1), corroborating previous evidence suggesting a link between aerobiosis and GC% in prokaryotes (13).

**Figure 2:**
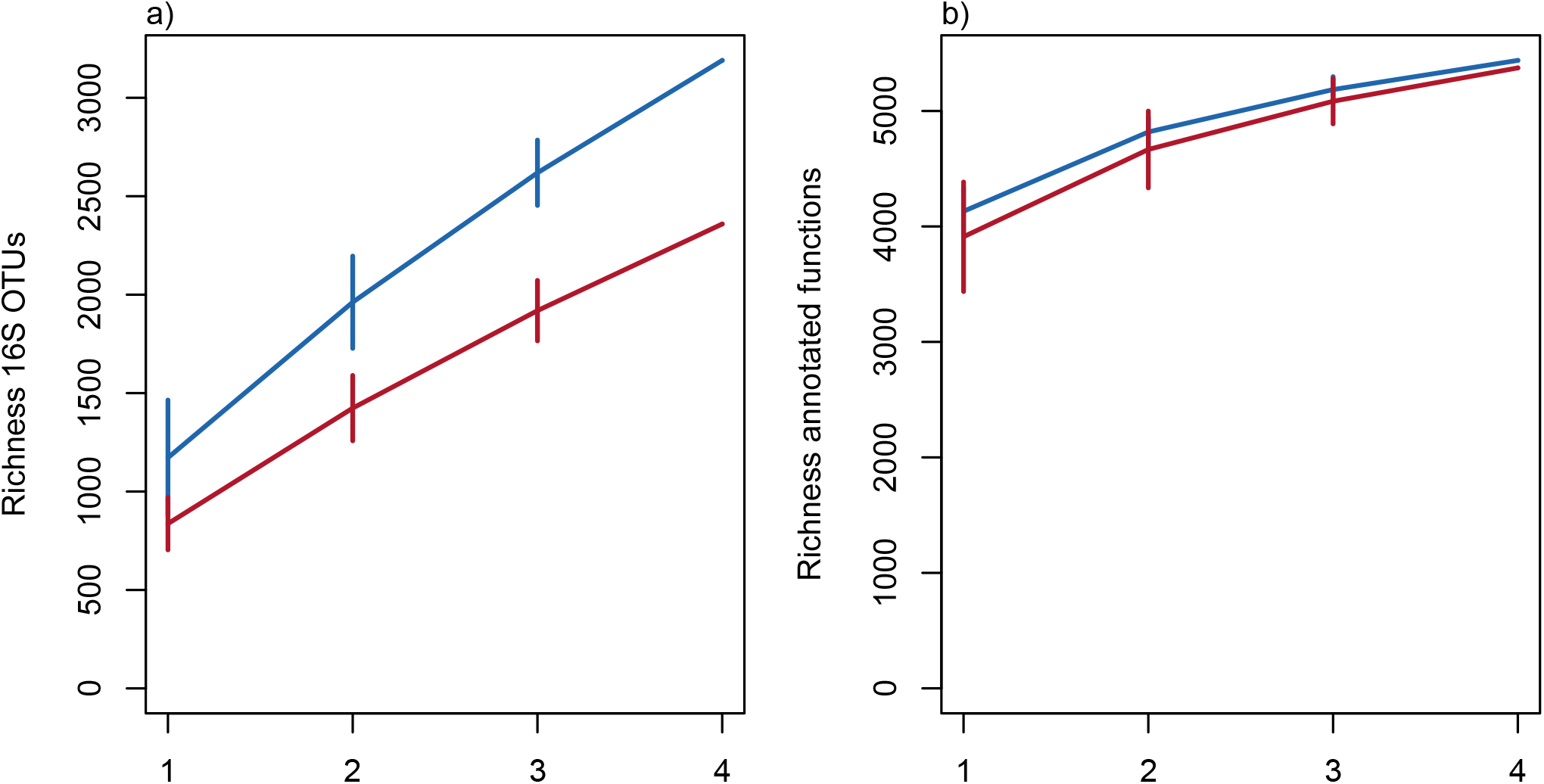
Within site and across site taxonomic and functional richness represented by site based accumulation curves. Standard deviations are calculated from random permutations of the data, with red lines representing low pH and blue lines high pH. (a) Taxonomic richness determined by 16S rRNA sequencing is higher at high pH, both within individual sites (alpha diversity) and accumulated across sites (gamma diversity); (b) richness of annotated functional genes following whole genome sequencing is only marginally lower in low pH at individual sites, and converges across all sites.

**Table 1:** Summary of soil and metagenomic characteristics. Functional annotations are available on MG-RAST under the following sample IDs.

| Sample | pH | Land use type | Organic matter (%) | Moisture (%) | No. reads | G:C content |
| --- | --- | --- | --- | --- | --- | --- |
| 4475877 | 4.5 | Heath | 21.2 | 56 | 400965 | 60 |
| 4475885 | 4.3 | Acid grassland | 70.4 | 82 | 612521 | 62 |
| 4475892 | 4.1 | Acid grassland | 47.8 | 72 | 647598 | 61 |
| 4475897 | 4.4 | Coniferous woodland | 88.8 | 90 | 645976 | 59 |
| 4475881 | 8.3 | Hedgerow | 24.1 | 48 | 556279 | 63 |
| 4475888 | 8.4 | Improved grassland | 6.4 | 25 | 621879 | 63 |
| 4475891 | 8 | Arable | 5.1 | 21 | 679130 | 64 |
| 4475893 | 8.5 | Improved grassland | 8.9 | 33 | 722025 | 63 |

Mean richness (alpha diversity) of functional genes was 4153 and 3915 in high and low pH soils, respectively (figure 2b); though this difference was not statistically significant (t-test, p=0.12). This was in contrast to the amplicon data, where richness was consistently higher across all samples at high pH (t-test, p<0.05). No correlative associations were observed between functional and taxonomic richness, in contrast to one previous study across a range of prairie grasslands (8). Importantly however, we found that whilst the accumulation of taxon richness over sites accentuated the differences in taxonomic diversity, this was not true for functional richness where 4 low pH soils possessed equivalent total functional diversity to 4 high pH soils. Thus, it is clear that whilst taxonomic diversity may be restricted by low pH related factors, functional diversity across multiple samples can be maintained through higher between-site variance (beta diversity) possibly mediated through enhanced metabolic versatility within acidophilic taxa.

### Large differences in relative gene abundance between low and high pH soils

Since a lack of difference in functional richness does not mean that soils are functionally similar across the pH gradient, we next sought to assess differences in relative abundances of functional genes. Figure 3 shows the overall abundance of genes classified at the broadest level (level 1 subsystems classification). Despite similarities in the abundance ranked order of hierarchically classified genes, a number of notable differences between soils of different pH were immediately apparent for several relevant functional categories, such as amino acid cycling, respiration, membrane transport, stress and virulence-disease-defense. However, a lack of difference at this level (notable for carbohydrate and nitrogen metabolism) could be due to significant differences within higher level functional categories. To address this, we performed a multivariate assessment of gene composition classified at the level of function. This revealed large differences between soils of different pH; indicating that the pH defined communities shared more similar functional gene profiles independent of their geographical origin (figure 4a, anosim p<0.05). The analyses based on relative abundance therefore shows that functional genes were differentially abundant in opposing soil pH classes; and in combination with higher functional beta diversity at low pH explains the equivalence of alpha diversity metrics.

**Figure 3:**
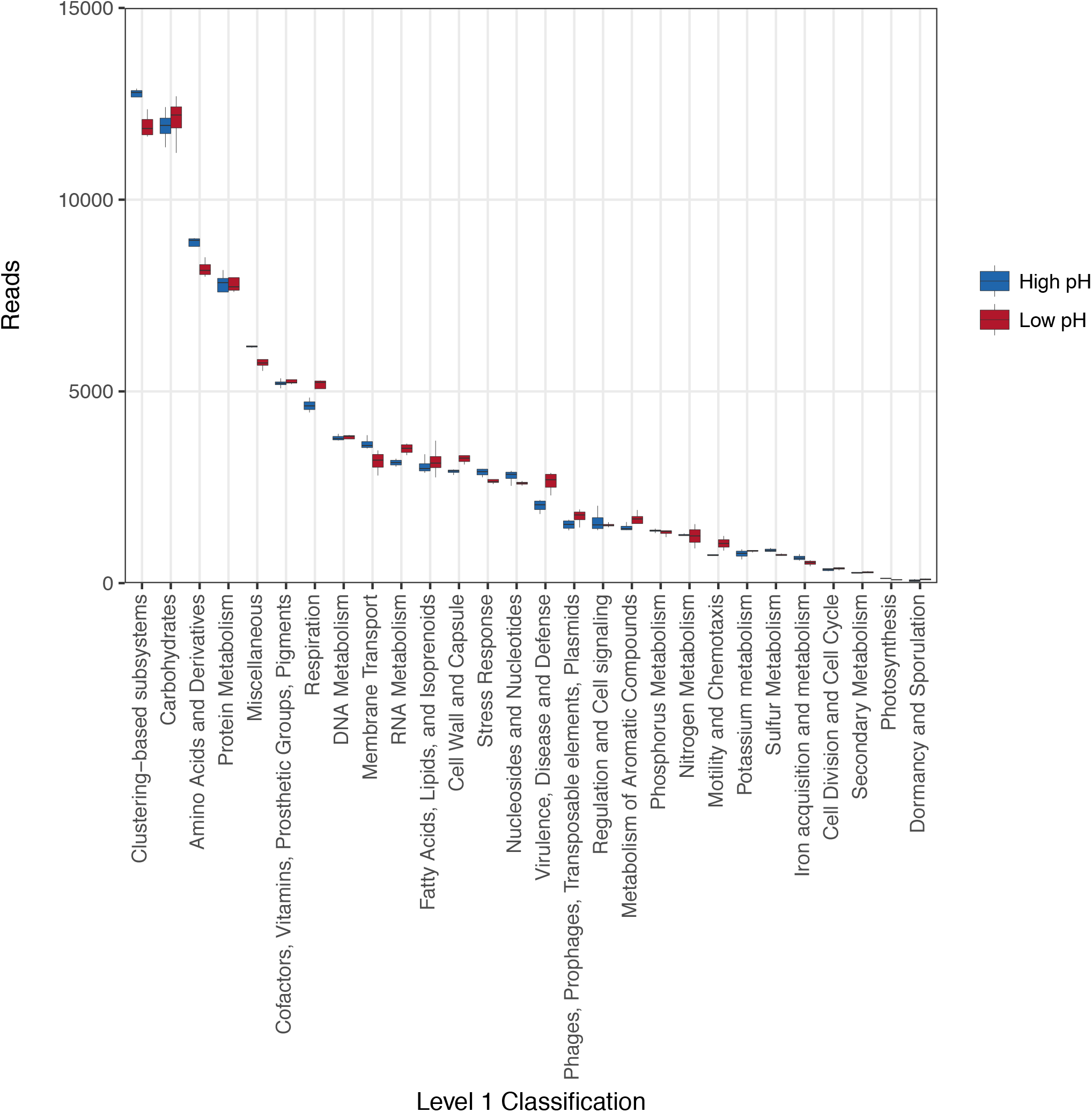
Abundances of annotated functional genes classified at the broadest level (level 1 subsystems classification), with total reads standardised across samples to 92442 reads.

**Figure 4:**
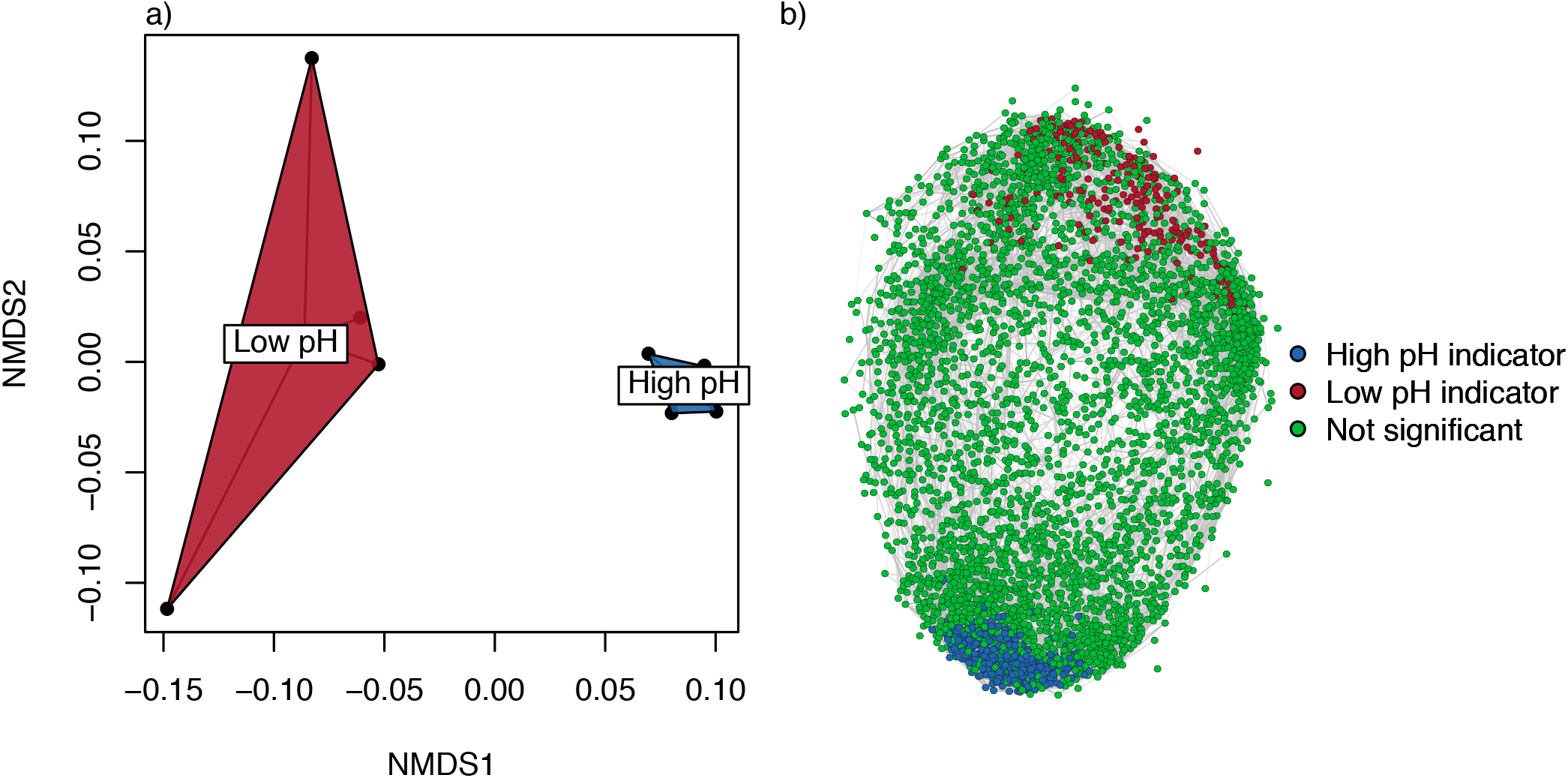
(a) Ordination of functional genes (classified at the level of function) using two dimensional NMDS reveals strong clustering of sites by pH irrespective of geographical sampling origin. (b) Network depicting strong (>0.9) positive correlations between annotated functional genes. For clarity, rare genes with less than 10 instances across all samples were omitted. Significant indicators are coloured according to pH class following indicator (indval) analyses.

Indicator gene analysis (Indval) was then used to define and explore the characteristic genes contributing to the differences between the pH defined soils. Of a total of 6194 annotated genes, 206 and 322 significant functional indicators were found for the low and high pH soils, respectively. A network depicting only strong positive correlations (>0.9) between genes across all samples revealed an expected lack of connectedness between opposing indicators, and in general low pH indicator genes were less correlated than high pH indicators reflecting the greater magnitude of functional beta diversity across low pH soils (figure 4b). To examine the functional identity of indicators, we constructed a circular plot (figure 5) at the functional gene level of classification, omitting rarer genes for ease of presentation (<50 reads across all samples), and labelling the indicators using the respective level 3 classification. A full table of all gene abundances, their classification and indicator designation are provided in the supplementary material (file S1). The following subsections discuss some relevant indicators of low and high pH soils. Whilst this discussion is by no means exhaustive, we divide the discussion into two sections based on physiological processes for survival and nutrient capture; and metabolism.

**Figure 5:**
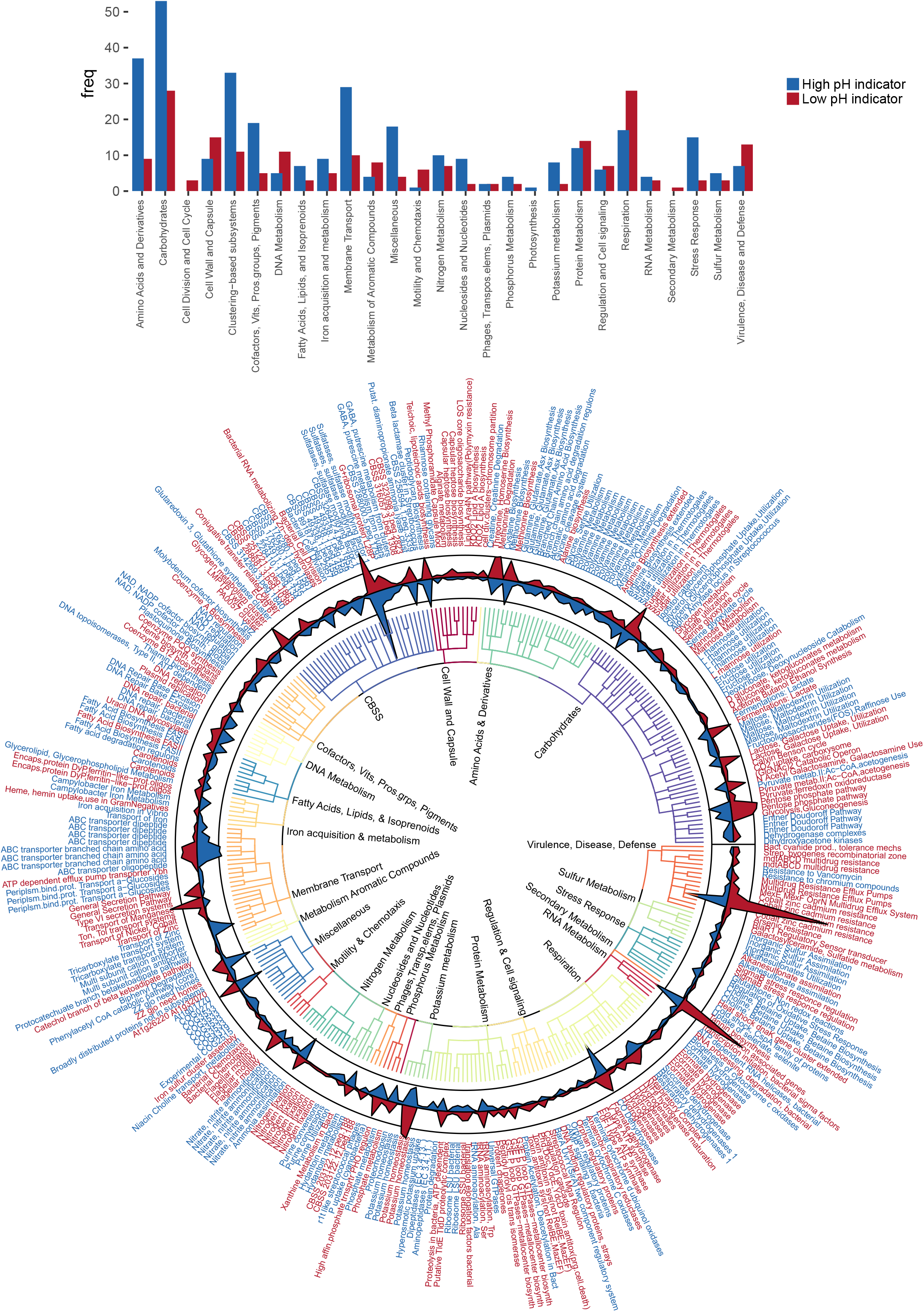
(a) Bar plot showing the frequency of indicator genes at the broad level 1 classification. (b) Circular plot displaying the identity and abundances of indicator genes for low and high pH soils. Nodes represent individual functional indicators, though are labelled with the more descriptive subsystems level 3 classification, i.e. repeated node labels indicate different functional indicators within the same level 3 subsystems classification. Node labels are coloured red and blue for particular genes that are significantly more abundant in low or high pH soils, respectively. Line plots represent total abundances of the indicators within the rarefied datasets, and are filled according to pH (red=low, blue=high). The tree depicts the hierarchical subsystems classification, with level 1 classifications being labelled on the internal nodes.

### Variable physiological strategies for survival and nutrient acquisition

The indicator analysis reveals an array of interlinked physiological adaptations to life at opposing ends of the soil environmental spectrum (figure 5). Firstly, with respect to cellular physiology, the high pH soils possessed a greater abundance of ABC transporters relatable to nutrient acquisition (14). In addition to the abundant amino acid, peptide and tricarboxylate transporter indicators (level 1 classification: membrane transport), numerous other transporters were significantly enriched at high pH though under different subsystems classifications. These included the majority of carbohydrate indicators for mono and disaccharide uptake, as well as other transporters for inorganic sulphate, cofactors, polyamines, ammonia/nitrate, potassium uptake proteins, and high affinity phosphate transporters (15). Interestingly, transporters for iron acquisition as well as osmoprotection were also evident at high pH, possibly reflecting low moisture and iron availability in the selected high pH soils. Together these findings indicate that the high pH soils can be distinguished functionally from low pH soils on the basis of a greater abundance of transporters for the direct uptake of available substrates and cofactors required for growth.

The membrane transport related indicators of low pH soils included only two low abundance genes related to nutrient acquisition (phosphate and glucose, not indicated in figure 5 due to low abundance), but a number of genes linked to metal acquisition (ybh, MntH, HoxN/ HupN/ NixA) and protein secretion (siderophores and extracellular enzymes). Relatedly a number of membrane proteins (mainly proton antiporters) for the efflux of antibiotics and toxic compounds were characteristic of low pH soils (ACR3, BlaR1/MecR1, CusA, CzcC, MerR, MacB, NodT, MdtB, MdtC), but annotated under ‘Virulence, Disease and Defense”. Indeed, the gene for cation efflux protein cusA related to cobalt-cadmium-zinc resistance was one of the more abundant genes across all metagenomes but was significantly enriched in low pH soils. Given the nature and location of the low pH soils examined it is unlikely these represent adaptations to either severe metal contamination or anthropogenic sources of antibiotics, but rather are related to pH enhanced reactivity of toxicants (e.g. metal ion solubility generally increases with decreasing pH) and the alleviation of acid stress through membrane efflux (16–20). It is also conceivable that they may be required for metal import for a variety of metal necessitating enzymes (see below) since many are proton/cation antiporters. Supporting this, another strong indicator of low pH was a potassium transporting ATPase gene (level 3 class: potassium homeostasis) coding for a membrane protein responsible for exchange of H^+^ and K^+^ ions across the plasma membrane, suggesting the coupling of acid stress response with elemental acquisition. As an aside, we were unable to find any studies that have examined acid soils for their potential as a source of antibiotic resistance, but our findings implicate adaptation to acidity or anaerobiosis as a possible factor underlying natural resistance to antibiotics (21, 22). Other notable indicators of low pH included chemotaxis and motility genes, plausible given the higher moisture contents of these soils (18). In total, these results identify that in the acidic soils considerable energy investments must be made in cellular processes to survive in an acid stressed, oxygen limited and low nutrient environment.

### Metabolic potential of contrasting pH soil communities

Carbohydrate processing was one of the most abundant broad classes of annotated functional processes and within this level 2 class serine-glyoxylate cycle, sugar utilization and TCA cycle genes were identified as the most abundant. Despite the expected conservation of many key metabolic functions, a number of notable indicators were found (figure 5). Several genes differed for the processing of mono and oligosaccharides, with a number of genes for L-rhamnose and fructose utilization being of greater abundance at high pH; whereas several extracellular enzyme coding genes for glucosidase, beta galactosidase, mannosidase and hexosamidase were elevated at low pH. With respect to the central carbon metabolism, certain components of the Entner-Doudoroff pathway were reduced in abundance in low pH soils (gluconate dehydratase, gluconolactonase); but by far the most abundant and significant low pH indicator was the xylulose 5-phosphate phosphoketolase gene. This gene along with another strong low pH indicator – transketolase – is part of the pentose phosphate pathway, a metabolic pathway parallel to glycolysis yielding pentose sugars and reducing agents. Though not previously considered largely with respect to soil functionality, this gene was generally of high abundance across all soils. However, novel position-specific isotope tracing experiments have recently provided functional evidence of the significance of the pentose phosphate pathway in soil C cycling (23). Its significance here in low pH organic soils remains to be questioned, although we note it is involved in fermentative processes and therefore would be expected to occur more frequently in these more anaerobic soils (24). Further evidence of an increase in fermentative processes in low pH soils was seen with a number of indicator genes for pyruvate:ferredoxin oxidoreductase, lactate fermentation (carbohydrate metabolism) and a wide range of hydrogenases (25, 26). Several recent studies have demonstrated that these hydrogenases are widespread in soil microorganisms (25, 27, 28), and they may also represent another means of consuming protons (29). Among these were cytoplasmic NAD-reducing hydrogenases that may be linked to respiration and fermentation through their NADH generation capabilities (30, 31). The nickel transporters (HoxN/HupN/NixA family), identified earlier may also be linked to this process since nickel is required for the metal center of NiFe hydrogenases (32, 33). Together, the metagenomic data suggest that low pH soil communities harbor adaptive physiological strategies of using molecular hydrogen oxidation and coupling it with respiration and fermentation to generate energy (34).

Whilst representing only a small proportion of these metagenomes, a number of nitrogen metabolism associated genes differed significantly between the soils of different pH. Nitrogen fixation genes were consistently strong indicators of low pH soils, with a number of genes found for molybdenum dependent nitrogenases (figure 5). Indeed, some of the nitrogenase indicators were unique in being universally present at low pH but entirely absent in the high pH soils. These findings indicate that microbial N input into soils either through symbiotic or non-symbiotic routes may be relatively larger in acidic soils, and there was evidence from the taxonomic assignments that the Bradyrhizobia may play a key role in this process (not shown). Low pH soils are characterized by decreased decomposition rates that could reduce the available N in soils necessitating microbial N fixation (35–37). The coupling of N fixation with abundant hydrogenases may also represents an efficient system for recycling the H_2_ produced in N_2_ fixation, minimizing the loss of energy (32, 38). High pH soils showed significantly more genes linked to nitrate/nitrite ammonification, ammonia assimilation, and denitrification (figure 5). The dominant pathways appeared to be related to ammonification (nitrate reduction to ammonia) and ammonia assimilation, and these were more abundant in high pH soils (39). Relatedly, a number of notable indicators of high pH soils were for the degradation of amino acids and derivatives, such as arginine, ornithine, polyamines, urea and creatine. Coupled with greater abundance of amino acid and peptide transporters, these observations infer an increased reliance on scavenging N enriched compounds originating from biotic inputs in high pH soils (20, 40, 41). Conversely, the lower abundance of these genes at low pH may indicate reduced bioavailability of these compounds possibly due to increased soil adsorption with greater cation exchange capacity (42).

## Conclusions

Our study shows that despite large differences in the taxonomic diversity of bacteria known to exist across soil environmental gradients, there was little evidence to suggest this results in large differences in the diversity of functional genes. Rather, low and high pH soils differed in the relative abundance of specific functional genes, and these indicator genes reflected differences in survival and nutrient acquisition strategies caused through adaptation to different environments. For the low pH soils, there were a number of abundant functional genes that highlight the importance of varied biochemical and physiological processes which we infer to be important adaptations to life in acidic, wet and oxygen limited environments. Indeed our results highlight the coupled action of acidity and anaerobiosis in mediating bacterial functional responses (22). In such soils a considerable investment of energy must be made into complex processes for capturing nutrients and energy, as well as stress responses caused by both the exterior environment and cellular metabolism. In combination, this is reflected by a greater abundance of cation efflux pumps, C and N acquiring systems such as direct fixation, and fermentation (figure 6). Higher pH soils conversely possessed more direct transporter mediated mechanisms for C substrate acquisition together with numerous indicators of organic N acquisition and consequent cycling (figure 6).

**Figure 6:**
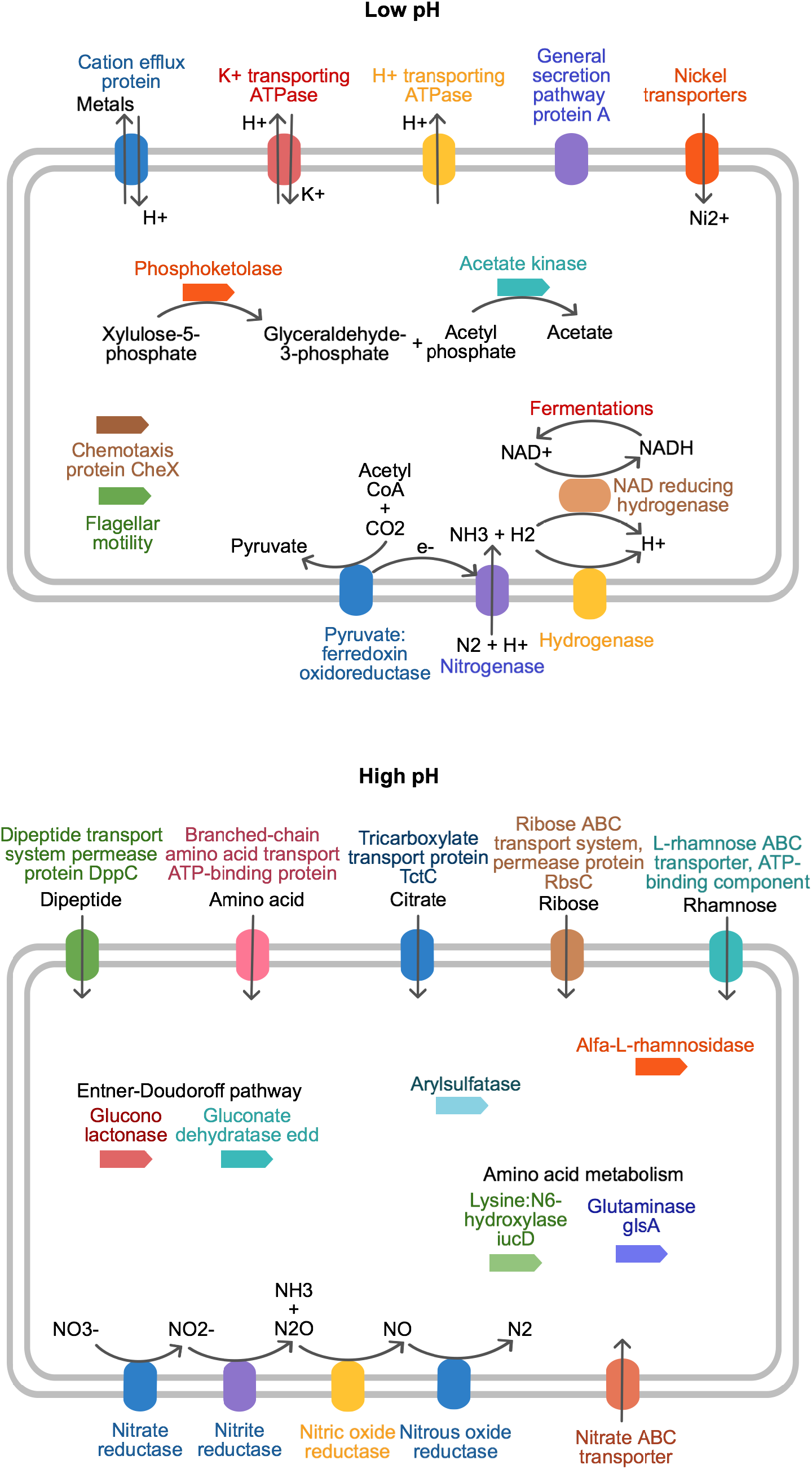
Schematic summarising some of the main physiological differences for survival, nutrient acquisitions and substrate metabolism across the pH gradient, as identified from the indicator analyses. We note that inclusion of a gene in either schematic is based on differences in abundances and does not implicate the presence/absence of a particular pathway across the gradient (refer to figure 5b).

In identifying specific indicators across the gradient, we acknowledge some limitations to the metagenomic approach. Firstly, the analyses are reliant on accurate functional assignment of reads, and there are recognized issues with respect to both bioinformatic annotation and the experimental assignment of a sequence to a single function (43). For this reason, we have focused our discussion on the more prevalent indicators represented by a variety of individual functional gene categories. An additional concern is that observed changes in relative gene abundance could be simply due to change in an unrelated gene (44). Whilst various methods for standardizing reads have been applied (such as relating abundances to rRNA genes, or calculating proportions within discrete subsystems) these approaches are not entirely without scrutiny. We prefer to consider the analyses akin to a mass balance – whereby the relative abundances of genes reflect the proportion of investment made for a given amount of nutrient into different proteins. The relevance for function at the ecosystem scale is a separate line of enquiry necessitating process measurement and assessment of biomass size, and we envisage metagenomic studies will provide more relevant functional targets.

In conclusion, we identify that considerable metabolic diversity and variability can exist within communities of environmentally constrained taxonomic diversity. Our intent at focusing analyses at broad ends of the pH spectrum was inspired to encompass an assessment at the extremes of soil functionality. In doing so, we make available sequence datasets which may be useful to others looking to assess the diversity of specific functional genes for a wide range of soil processes across a range of soils. Furthermore, we envisage that the indicators identified here, whilst being from an extreme soil contrast, may also be relevant at local scales for understanding more subtle alterations in soil function. For example, more “natural” soils in temperate climates typically store more carbon, tend towards acidity, and have increased moisture retention; whereas human agriculture forces soils to neutrality and depletes soil carbon and moisture. Our results may therefore be relevant in understanding the balance of energy and C storage mechanisms under altered land management; as well as permitting future design of smarter systems for efficient soil nutrient capture and recycling.

## Materials and Methods

Characteristics of soil examined and location of sampling sites are shown in Table 1 and Fig. 1, and full details of the sampling, nucleic acid extraction and taxonomic analyses are provided in our previous manuscripts (2, 7). Eight soils were selected for detailed metagenomic analysis on the basis of soil pH alone and are representative of the extremes of both the soil environmental conditions and bacterial diversity range encountered across Britain. Bacterial communities were characterized using tagged amplicon sequencing as previously described (7) using primers 28F / 519R, and sequencing on the 454 pyrosequencing platform through a commercial provider. Raw sequences from amplicon sequencing were analyzed using QIIME, using UCLUST to generate OTU’s at 97% sequence similarity. For whole genome metagenomic sequencing, three μg of total DNA from each sample was submitted to the NERC Biomolecular Analyses Facility (Liverpool, UK) with two samples per run on the 454 platform (sequencing statistics in Table 1). Resulting sequences from metagenomic analysis were annotated with the Metagenomics Rapid Annotation using Subsystems Technology (MG-RAST) server version 4.0 (45). Functional classification was performed using the SEED Subsystems database with a maximum e-value cut-off of 10^-5^, minimum identity cut-off of 60% and minimum length of sequence alignment of 15 nucleotides. Gene abundance tables derived from MG-RAST were imported into R for downstream analyses. Rarefaction, species diversity accumulation, ordinations, and statistical analyses were performed using the vegan package (46) under the R environment software 2.14.0 (R Development Core Team 2011).

